# Spontaneous and stimulus-driven arousal produce distinct acetylcholine dynamics across sensory and prefrontal cortex

**DOI:** 10.64898/2026.06.02.729441

**Authors:** Anna R. Chambers, Eyal Y. Kimchi, Yurika Watanabe, Tatenda Chakoma, Daniel B. Polley

**Author notes:** Department of Neurology, Northwestern University, Chicago IL 60208. These authors contributed equally (Anna R. Chambers, Eyal Y. Kimchi).

## Abstract

Acetylcholine (ACh) release from the basal forebrain has traditionally been viewed as a slow, spatially diffuse signal regulating cortical arousal across sleep and wakefulness^1–3^. Recent characterizations with higher resolution optical sensors have revealed rapid, local cholinergic modulation supporting dynamic changes in sensory processing, associative learning, and behavioral state^4–7^. However, sensory events that recruit cortical ACh often also change arousal and evoke movements, making it difficult to determine whether ACh transients reflect sensory features of environmental stimuli or the behavioral state changes that accompany sensory stimulation. To separate these contributions, we performed optic fiber recordings of a genetically encoded ACh fluorescent sensor in the auditory, visual, and prefrontal cortex of awake, head-fixed mice while monitoring pupil size and facial movements. Across cortical areas, ACh release tracked spontaneous fluctuations in arousal state, as indexed by pupil dilation and orofacial movements. Sensory stimuli also evoked rapid ACh transients, with sounds producing larger and more widespread responses than visual stimuli. Because sounds also elicited time-locked pupil dilations and facial movements, we used multivariate modeling to estimate the relative contributions of stimulus features, arousal, and behavior to cortical ACh dynamics. We identified a regional dissociation: sound-evoked ACh release in prefrontal cortex was largely explained by arousal- and movement-related variables, whereas auditory cortical ACh release retained a stronger relationship to stimulus features. These findings show that cortical ACh signaling reflects both shared arousal state and area-specific sensory processing and demonstrate that sound is especially effective at recruiting rapid, widespread cholinergic modulation across cortex.

## Introduction

Neural recordings from awake, behaving animals have revealed that cortical responses are dominated by widespread, low-dimensional signals associated with brain and behavioral state^8,9^. Recent studies suggest that uninstructed behavioral responses, such as facial movements, may indirectly account for a large degree, though not all^10^, of spatially distributed cortical spiking previously attributed to cross-modal sensory tuning^9,11^. The traditional picture of cortical neuron activity reflecting functional topography and sensory receptive field tuning now exists alongside an updated version, where even primary sensory cortices are largely readouts of internal brain state and overt behaviors^5^.

To what extent does a cortical response reflect the sensory features of the stimulus, rather than the indirect recruitment of involuntary behavioral responses and arousal-related neuromodulator release? The ‘direct sensory’ versus ‘indirect behavioral’ dichotomy is difficulty to distinguish; notably, cortical networks receive projections from neuromodulatory centers that signal arousal and behavior^12^, yet neuromodulatory nuclei themselves receive direct sensory inputs^13–16^. Cholinergic projections to the cortex originating from the basal forebrain (BF) show modality-specific topography^17^ and stimulus feature specificity, even to passively presented stimuli^13,17–19^. At the same time, cholinergic neuron activity is associated with spontaneous arousal and facial movements in the absence of stimuli^12,20^. Thus, a single upstream source could carry information about external events and their behavioral impact in parallel, ultimately informing the rapid detection of potential threats and the selection of appropriate responses.

The unique influence of the stimulus on cortical acetylcholine (ACh)—and subsequently, local neural activity and plasticity— is not necessarily a question of ‘whether,’ but of ‘what’ and ‘where.’ However, the answers are particularly difficult to parse for stimuli that elicit robust arousal and non-habituating behavioral reactions, such as loud, broadband sounds^10,11,21,22^.

In this study, we explored the stimulus specificity, behavioral dependence, and spatial topography of cortical ACh release with fiber photometry and quantitative videography. We analyzed the association of cortical ACh release to external metrics of brain and behavior state—pupil diameter and orofacial movements. To disentangle the unique contributions of stimulus, behavioral response, and brain state to local ACh release, we varied the modality, complexity, and intensity of stimuli to elicit varying combinations of endogenous and exogenous arousal responses. We then compared the contributions of each element across sensory and non-sensory cortical regions. We highlight a potentially privileged role for complex sounds in eliciting stimulus-dependent ACh release across auditory and visual sensory regions, while ACh in the prefrontal cortex, a non-sensory association area, appears to primarily reflect behavioral indicators of internal arousal state.

On a broader level, these results provide a narrative for how and why certain types of sensory stimuli—for example, those that are more likely to signal dangers—may elicit particularly widespread and rapid behavioral and computational shifts, coordinated across distributed cortical networks^23,24^.

## Results

### Movement- and pupil-based readouts of arousal are associated with different degrees of ACh release in sensory and non-sensory cortical regions

We first analyzed the relationships between cortical ACh, pupil dilation, and orofacial movements in the absence of sensory stimuli. We performed adeno-associated virus-mediated delivery of a fluorescent ACh indicator, GRAB_ACh_,^7^ to the visual, auditory, or prefrontal cortex of adult mice (**Fig 1A**; ACtx N = 15; VCtx N = 8; PFC N = 7) and combined fiber photometry recordings of ACh release with continuous facial video monitoring while mice were head-fixed and sat in a tube. The pupil was identified across frames with DeepLabCut^25^, while orofacial motion energy (FME) was extracted from the video pixels encompassing the cheek^21^ (**Fig 1B**).

**FIGURE 1.**
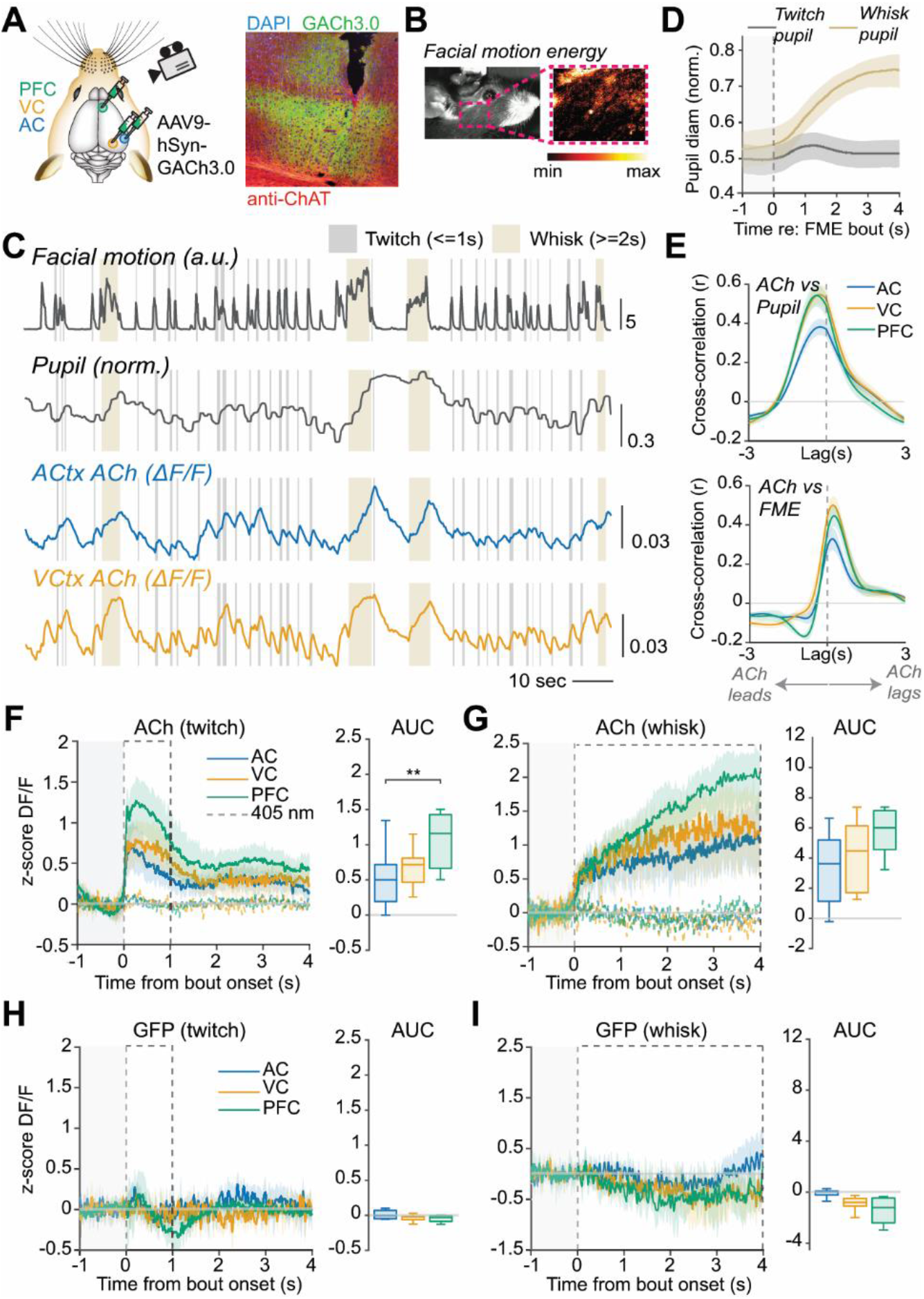
(**A**) Schematic depicting the locations of AAV injections delivering GRAB_ACh_ to the PFC, VCtx, and/or ACtx in the mouse cortex (right), and histological section from the cortex showing GRAB_ACh_ and cholinergic terminals identified with immunostaining. Profile of implanted tapered fiber used for photometry visible in section. (**B**) Single frame from a video of a head-fixed mouse’s face, with the location of analysis for facial motion energy (FME) noted within the pink dotted rectangle. Zoomed in version shows calculated FME in the identified pixels across frames. (**C**) Representative recordings of FME, pupil diameter, and ACh (simultaneously measured from two separate fibers implanted in ACtx and VCtx) in a passive, head-fixed mouse in the absence of stimuli. Gray and beige boxes identify twitch (significant FME bout <= 1s in duration) and whisk (FME bouts >=2s in duration) events, respectively. (**D**) Normalized pupil diameter aligned to the onset of twitch or whisk bouts (mean +/- s.e.m.). (**E**) Cross-correlation between cortical ACh (in ACtx, VCtx, or PFC) and pupil (top) and FME (bottom). (**F-G**) Z-scored DF/F of ACh signal (measured at 465nm) in each cortical region in response to twitch (**F**) or whisk (**G**) events, compared to the control signal measured at 405nm (dotted lines). Box plots show the median +/- inter-quartile range of the ACh signal, quantified as the area under the curve in the timeframe 0-1s (twitch) or 0-4s (whisk). ACtx showed significantly less ACh release during spontaneous facial twitches compared to PFC (Wilcoxon rank-sum test; *p* = 0.0036). (**H-I**) Same as (**F-G**), but in mice with AAV-mediated delivery of only the fluorophore GFP.

Pupil dilation and facial movement are often used as behavioral proxies of arousal state in mice^5,12,26^. We found that pupil size, facial movement, and ACh levels all fluctuated spontaneously on time scales in the range of 0.1 – 1Hz and were broadly coherent (**Fig 1C**). Most of these fluctuations corresponded to brief twitches of the face and associated phasic ACh, however occasional bouts of whisking were accompanied by more sustained ACh release (**Fig 1D**). Though pupil, FME and ACh were largely correlated, they unfolded with distinct temporal dynamics (**Fig E**), with ACh signals leading pupil dilations and following FME bouts.

ACh levels increased sharply after smaller, shorter-duration facial “twitches” (**Fig 1F**) and larger, longer duration bouts of whisking (**Fig 1G**). A clear correspondence between ACh release and facial movements was observed in all three brain regions, though PFC displayed significantly higher levels of ACh release than ACtx, with VCtx in between (**Fig 1F-G, right**).

To identify the contribution of artifacts in these fluorescence measurements (e.g. intrinsic signals or brain tissue movement), we analyzed two control conditions: First, the fluorescence signal measured from the control excitation wavelength (405 nm, **Fig 1F-G**, dashed lines); Second, the fluorescence signal measured from a separate cohort of mice expressing a control GFP fluorophore rather than the GRAB sensor (ACtx N = 7; VCtx N = 7; PFC N = 4; **Fig 1H-I**). The twitch- and whisk-aligned fluorescence signals seen in the GRAB_ACh_ mice were absent in both of these control conditions.

### Auditory gratings are more effective than visual gratings at eliciting ACh release in sensory and non-sensory cortical regions

Though the basal forebrain is closely associated with arousal recruitment, multiple studies have confirmed that cholinergic and non-cholinergic neurons of the BF respond to sensory stimuli^15– 17,27^. Therefore, we investigated the impact of visual and auditory stimuli on ACh release across cortical regions. We chose to compare complex, time-varying stimuli across two modalities: sinusoidal visual gratings, and auditory gratings (also known as auditory ripple stimuli^28^, in which the sound amplitude envelope oscillates sinusoidally across the spectral and temporal domains). Experiments in GFP-expressing control mice demonstrated the contribution of non-specific intrinsic signal changes at longer latencies, prompting us to focus only on the immediate post-stimulus period, from 0 to 1 sec after stimulus onset in GRAB_ACh_ mice (**Fig S1**). The 405nm control wavelength was less sensitive to sensory-evoked intrinsic signal contributions (**Fig S2**).

We found that auditory gratings elicited robust ACh release in ACtx, VCtx and PFC, even at moderate or low intensities (**Fig 2A-C**). ACh release elicited by visual gratings was weaker overall than auditory gratings. Responses to the highest contrast stimulus were prominent in the ACtx and PFC, but moderate- and low-contrast stimuli did not elicit significant ACh responses in any cortical region (**Fig 2D-F**).

**FIGURE 2.**
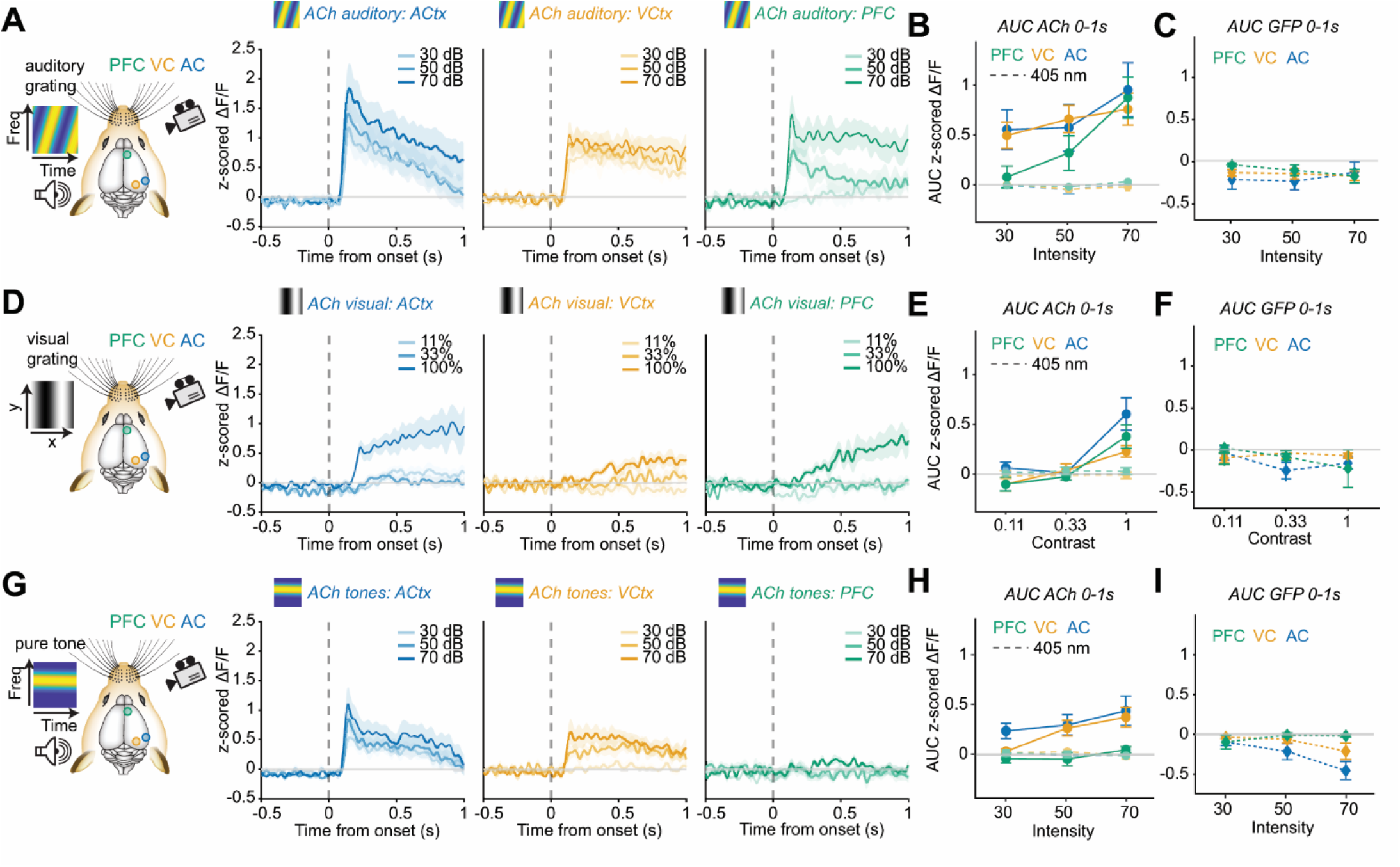
(**A**) ACh release measured in ACtx, VCtx, or PFC in response to auditory gratings of varying intensities. Traces show mean +/- s.e.m. (**B**) Mean ACh signal in the 0-1s timeframe, quantified as area under the curve, for all intensities, compared to the 405nm control (dotted lines) or fluorescence from the GFP mice (**C**). (**D-I**) Same as (**A-C**), but in response to visual gratings (**D-F**) or pure tones (**G-I**).

Though complex sounds appeared to be more effective than comparable visual stimuli at eliciting ACh release across the cortex, it is impossible to directly compare the stimulus effect across two different modalities. To further dissect the impact of sounds on cortical ACh release, therefore, we introduced the most rudimentary sound stimulus: pure tones (**Fig 2G-I**). Pure tone stimuli elicited ACh release in ACtx and VCtx—but not PFC— even at moderate intensities (**Fig 2H**). **Table S1** provides a compendium of evoked responses in individual mice to all stimulus types and intensities.

### Auditory gratings elicit significant pupil dilation and facial movements, which are correlated with cortical ACh release

Previous studies have shown that sensory stimuli—particularly loud, broadband auditory stimuli—induce stereotyped orofacial motor responses and pupil dilations^10,11,21,29^, underscoring that the behavioral indices of spontaneous arousal fluctuations can also be recruited by sensory stimuli. To investigate this relationship in greater detail, we quantified the arousal and motor recruitment of each stimulus type and intensity (**Fig 3**). Auditory gratings at moderate and high intensities elicited robust pupil dilation and facial movement, while visual gratings did not elicit significant behavioral responses even at the highest contrast (**Fig 3A,B**), in keeping with recent reports^10,21^. Meanwhile, pure tone stimuli elicited significantly lower behaviorally indexed arousal responses compared to complex sounds (**Fig 3C,D**; beige traces show auditory grating response averaged across intensities), and reached significance only for the highest intensity **(Fig 3C**).

**FIGURE 3.**
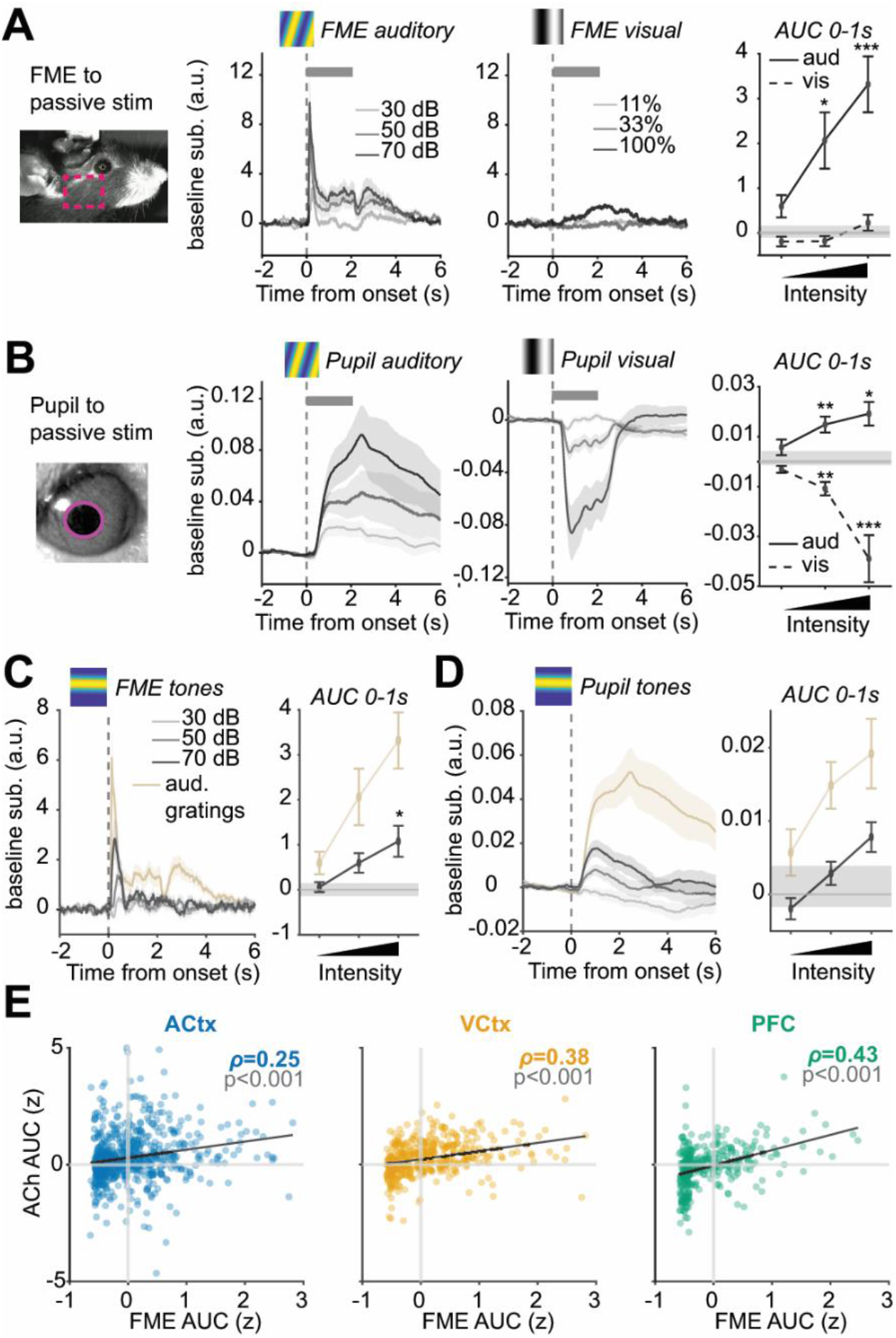
(**A**) FME in response to auditory or visual gratings; quantification at left shows the mean +/- s.e.m. of FME in the 0-1s post-stimulus period, across intensity/contrast. Gray horizontal bar denotes noise floor of FME calculated from silent/no stimulus trials, while asterisks denote stimuli that elicited significantly more FME than the noise floor (one-sided *t*-test). (**B**), Pupil diameter in response to auditory or visual gratings, quantified as in (**A**). Note pupil constriction to visual stimuli reflecting luminance changes rather than arousal changes. (**C-D**) FME and pupil diameter in response to pure tones at different intensities, with average auditory grating response shown in beige for comparison. (**E**) Scatter plots showing the correlation between FME and cortical ACh, which was significant in all cortical regions and most prominent in PFC.

While stimulus-driven ACh release was most strongly correlated with concurrent orofacial motion in PFC, agreeing with the results from spontaneous movements in silence (**Fig 1**), cortical ACh showed a significant positive correlation with orofacial motion in all regions (**Fig 3E**). Taken together, orofacial movements and pupil-indexed arousal were not obvious predictors of ACh release for pure tone and visual grating stimuli—robust ACh release was measured in the cortex even for stimuli that did not elicit significant movement or pupil dilation. However, the picture was less clear for auditory gratings, which elicited large degrees of movement, pupil dilation, and ACh release in all cortical regions.

### ACh release explained uniquely by the auditory grating stimulus is highest in ACtx, but still present in VCtx and PFC

ACh release to auditory and visual stimuli could be due to direct stimulus effects, behavioral and brain state context, or a combination of the two. To disentangle these possibilities, we analyzed the trial-by-trial differences in behavior, brain state, and stimuli to analyze unique and shared predictors of ACh variability (**Fig 4**). First, we identified brief orofacial twitches in the spontaneous (no stimulus) condition, and amplitude-matched FME bouts that were aligned with stimulus trials (**Fig 4A-C**). We then analyzed the concurrent ACh signals in the three cortical regions to ask whether a spontaneous FME bout would elicit comparable ACh release to an FME bout of the same size, co-occurring with a sensory stimulus. In ACtx, ACh traces were larger in the stimulus-evoked case, even while orofacial motion was matched across trials (**Fig 4D**). Meanwhile, ACh release was comparable, or even lower, for stimulus-evoked FME bouts compared to spontaneous bouts in VCtx and PFC, respectively (**Fig 4E-F**). This suggests that especially in ACtx, ACh release in the presence of a stimulus is significantly higher than what would be predicted by the ACh release associated with uninstructed behaviors.

**FIGURE 4.**
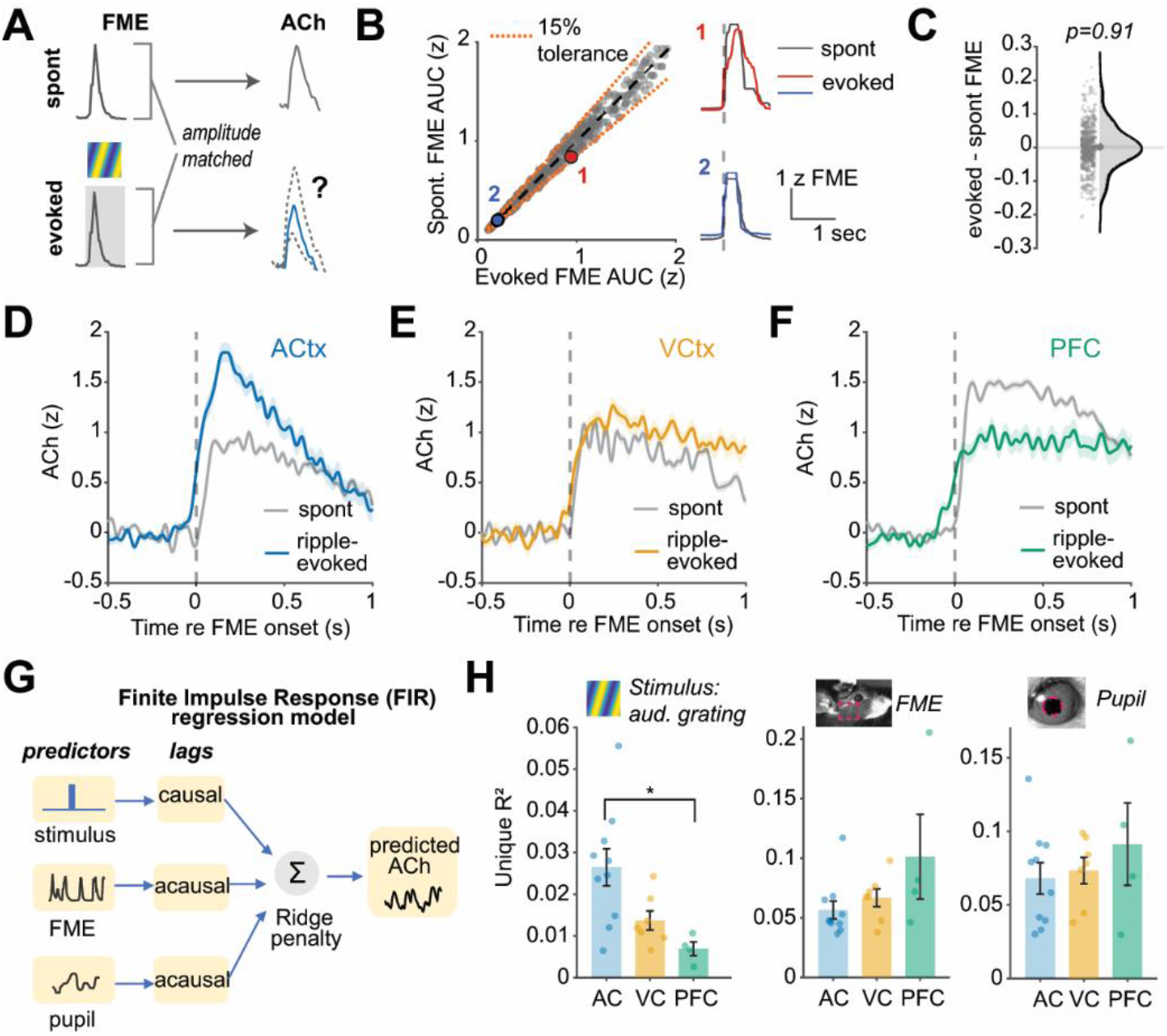
(**A**) Schematic showing example spontaneous FME bout (top right) and associated ACh transient (top left), compared with an amplitude-matched FME bout co-occurring with an auditory grating stimulus (bottom right). (**B**) Scatter plot showing the distribution of evoked and spontaneous FME bout sizes, quantified as area under the curve. Dotted red lines show 15% tolerance boundaries for amplitude matching. Numbered insets show example traces for matched spontaneous and stimulus evoked FME bouts. (**C**) Distribution of the size differences (area under the curve) between amplitude-matched spontaneous and stimulus evoked FME bouts. The mean of the distribution was not significantly different from zero (one-sided *t*-test). (**D-F**) Corresponding ACh signal during spontaneous (gray traces) versus amplitude matched auditory grating-evoked (colored traces) in each cortical region. Traces are mean +/- s.e.m. (**G**) Schematic of FIR regression model, with stimulus, FME, and pupil diameter as predictors of the ACh signal. (**H**) Variance partitioning analysis based on results from FIR regression model. Bar plots show the unique variance explained by each predictor in each brain region. The auditory grating stimulus showed the most unique variance explained in ACtx, significantly more than PFC (unpaired Wilcoxon signed rank test between cortical regions, *p* = 0.009).

To further disambiguate stimulus-dependent contributions to cortical ACh release, we employed a finite impulse response (FIR) regression model (**Fig 4G**), an extension of a multiple linear regression model. The model takes advantage of the fact that although pupil, facial motion, and ACh are positively correlated, they unfold on distinct timescales. When comparing sources of unique variability in the ACh release across cortical regions, ACtx, significantly more so than PFC, showed the most dependence on the stimulus (**Fig 4H**, left). Meanwhile, the unique variability due to the stimulus in VCtx was in between ACtx and PFC. FME and pupil diameter predictors trended in the opposite direction across brain regions (**Fig 4H**, middle and right), though this difference was not significant.

Overall, we noted that the auditory modality elicited a greater degree of ACh release overall. In sensory cortex, this ACh release could be attributed in part to the sensory stimulus itself, while ACh release in a non-sensory region, the PFC, was predominantly attributable to arousal and uninstructed behaviors. These spatially heterogenous features of the cortical ACh system reflects the topography of neuromodulatory projections from the brainstem, as well as the unique computations performed across diverse cortical circuits.

## Discussion

In this study, we used quantitative videography and fiber photometry imaging of a genetically encoded ACh indicator to analyze the relationships between sensory stimulus features, behavior, arousal, and spatially heterogeneous ACh release in the cortex. We found that complex sounds elicited robust pupil-indexed arousal, orofacial movements, and phasic ACh release. We sought to clarify the coupling between these behavioral and neurochemical metrics of arousal by varying stimulus properties such as modality, complexity, and salience; visual gratings and pure tones at moderate intensities did not elicit orofacial movements, however in ACtx and VCtx, ACh release persisted, suggesting a stimulus-dependent mechanism. Meanwhile, PFC exhibited some ACh release to visual gratings but none to pure tones. Analysis with a multiple linear regression model revealed that while the sensory stimulus accounted for some degree of unique ACh release variability in all three cortical regions, the effect was largest in ACtx. This feature of ACtx may support the ability of movement-sensitive neurons to accurately distinguish spontaneous from sound-evoked twitches in their spiking activity^29^.

Cortical responses are often attributed to ‘sensory’ or ‘behavioral’ sources, however it has long been acknowledged that rather than carried along separate parallel channels, the two types of information are necessarily integrated at early stages of brain processing^30^. This is true not only in the BF cholinergic system, but in other brainstem arousal control networks such as the lateral hypothalamus^31,32^, locus coeruleus^33^, and raphe nuclei^34,35^. Further, the interdependencies among stimuli, movements, and arousal are bidirectional: stimuli can elicit movements and arousal shifts, but sensory processing and sensory-guided behaviors reflect the pre-stimulus arousal and movement state^5,36^. In practice, it is difficult to disentangle the unique computational impact of a stimulus-evoked behavior, compared to a spontaneous expression of nearly the same behavior. The distinction, however, is important: neural circuits must unambiguously signal external, versus internal, triggers for arousal and motor recruitment to direct future decision making. Sounds are particularly useful external triggers, given the auditory system’s exquisite temporal precision and sensitivity across arousal states^37–39^.

The privileged role of the auditory modality for recruiting arousal and ACh release is not only reflected in the ethological value of sounds, but in the anatomy and physiology of the ACh system. In addition to its roles in arousal control and gating of plasticity, the basal forebrain also performs rapid high-resolution processing of auditory stimuli^15^. Cholinergic neurons’ responses to broadband sounds show latencies of only about 11 ms^16^, matching or even leading sound-evoked latencies in ACtx units, and their pure tone frequency tuning properties resemble those of neurons along the primary auditory pathway^15,16^. Short-latency sound information is routed to the basal forebrain via the medial subdivisions of the auditory thalamus^13,15,40^. These projections target the caudal tail of the basal forebrain, which is the main source of BF projections to the auditory cortex^13,19,41^. By contrast, PFC displays strong reciprocal connectivity with a more rostral BF region, the horizontal limb of the diagonal band of Broca (HDB), also the primary source of BF cholinergic inputs to VCtx^41^. The projections originating from HDB, rather than being strongly driven by sound, are more reliable readouts of behavior, arousal state, and reinforcement^17^. Taken together, the spatial heterogeneity we observed in sound-evoked ACh release mirrored the sound-encoding properties, or lack thereof, of a majority of that region’s BF projections.

The term ‘arousal’ can refer to a behavioral, physiological, or neurochemical state; its precise meaning in the context of neuroscience research is often unclear. As markers of arousal, orofacial movement and pupil diameter are straightforward to measure and analyze, but complicated to interpret: they are proxies of internal brain dynamics governed by multiple neurotransmitter systems^42,43^. Here, we measured these markers alongside cortical ACh, a sensitive readout that provides specific neurochemical context to the stimulus-evoked behavior and pupil changes. One caveat of the technique, however, is that ACh release could have originated from multiple sources, e.g. cholinergic projections originating from the BF or from the ventral pallidum^44,45^, or local ACh-releasing interneurons^46^. However, previous studies utilizing direct electrophysiological recordings from identified ACtx-projecting BF cholinergic neurons, as well as GCaMP imaging in the axons of these neurons in cortex, have shown that they are sound-responsive^15,17,19,27^. Because sounds are more associated with orofacial movement than visual stimuli^10,22^, the stimulus dependence of sound-evoked cortical ACh is more difficult to parse. Therefore, we used a linear model to ask whether trial-by-trial variability in the cortical ACh signal could be uniquely explained by the stimulus. To clarify the sensory circuits that drive stimulus-evoked ACh release, future studies could causally interfere, e.g. with optogenetics, with the neural pathways conveying sound or visual information from the periphery to central networks such as the cortex and BF. Finally, the current study is limited in its interpretation across behavioral contexts, as mice were head-fixed in our experiments (though sound-driven orofacial movement has been analyzed in freely moving mice^47^).

In addition to its causal role in brain state transitions^48^, the BF plays diverse roles in modulating neural activity and plasticity in cortical circuits^49,50^. BF cholinergic inputs to sensory cortex enhance stimulus responses and desynchronize cortical spiking^51–53^, modulation that has been posited to improve sensory perception^54,55^. Sensitive to arousal state, behavior, and sensory context, cortical ACh can thus accomplish multiple parallel operations to enable the detection and evaluation of environmental stimuli that could be critical for survival.

## Acknowledgements

We thank Yulong Li for making the GRAB_ACh_3.0 sensor available for purchase. Financial support was provided to DBP by NIH grant DC017078. EYK was supported by NIH grant MH116135.

## Author contributions

Conceptualization, A.R.C, E.Y.K. D.B.P.; Methodology, A.R.C., Y.W., T.C., E.Y.K. D.B.P.; Investigation, Y.W., T.C., E.Y.K.; Software, A.R.C. and E.Y.K.; Formal Analysis, A.R.C. and E.Y.K.; Data Curation, A.R.C.; Visualization, A.R.C, E.Y.K., D.B.P.; Writing – Original Draft, A.R.C. and D.B.P.; Writing – Review & Editing, A.R.C., E.Y.K, and D.B.P.; Resources, D.B.P.; Supervision, D.B.P.; Funding Acquisition, D.B.P.

## STAR ⋆ Methods

### EXPERIMENTAL MODEL AND SUBJECT DETAILS

All procedures were approved by the Massachusetts Eye and Ear Animal Care and Use Committee and followed the guidelines established by the National Institutes of Health for the care and use of laboratory animals. Experiments were performed in adult mice of both sexes, at least 2–3 months of age at the time the first measurement was performed. Male ChAT-cre-ΔNeo (homozygous, Jackson Labs 031661) and female Ai148 mice (hemizygous, Jackson Labs 030328) were bred in-house to generate mice of both sexes for a previous study^17^. Offspring used for the current study were hemizygous for ChAT-cre-ΔNeo and did not express GCaMP (ChAT+/GCaMP-). Offspring genotypes were confirmed by PCR (Transnetyx probes) and by histology following perfusion.

Prior to behavioral testing, mice were maintained on a 12 hr light/12 hr dark cycle with ad libitum access to food and water. Mice were grouped-housed unless they had undergone a major survival surgery. Fiber imaging of GRAB_ACh_3.0 sensor fluorescence was performed in 19 total mice, with each mouse contributing 1 or 2 brain regions (breakdown by cortical region as follows: 15 (ACtx), 8 (VCtx), and 7 (PFC) mice). A separate cohort of mice received AAV delivery of the fluorophore GFP for control experiments (N = 7 (ACtx); 7 (VCtx); 4 (PFC)).

## METHOD DETAILS

### Surgery, virus and head-fixation

Mice were anesthetized with isoflurane in oxygen (4% induction, 1.5-2% maintenance) and placed in a stereotaxic frame (Kopf Model 1900). A homeothermic blanket system was used to maintain body temperature at 36.6° (FHC). Lidocaine hydrochloride was administered subcutaneously to numb the scalp. The dorsal surface of the scalp was retracted and the periosteum was removed. The skull overlying the right cortex was exposed (for ACtx, by moving the temporalis muscle), and a burr hole was made using a 31-gauge needle. A motorized injection system (Stoelting) was used to inject 200 nL of AAV9-hSyn-ACh3.0 (diluted 10% in sterile saline from 3.45 × 10^13^ genome copies/mL) via a pulled glass micropipette 0.5 mm below the pial surface. We waited at least 10 min following the injection before withdrawing the micropipette. A tapered fiber (Optogenix, NA 0.39, diameter 200 µm, active length 1.0 mm) was implanted 1 mm below the pial surface and secured using dental cement dyed with India Ink, which also secured the titanium head plate. The exposed skull surface was prepped with etchant (C&B metabond) and 70% ethanol before affixing a titanium head plate (iMaterialise) to the skull with dental cement (C&B Metabond). At the conclusion of the procedure, buprenorphine (0.05 mg/kg) and meloxicam (0.1 mg/kg) were administered and the animal was transferred to a warmed recovery chamber. Sensor photometry experiments began 3 weeks following the injection.

### Video monitoring

Mice were placed in an electrically conductive cradle and habituated to head fixation during three sessions of 15 to 30 min over 3 consecutive days. Video recordings under isoluminous conditions were made at 30 Hz with a CMOS camera (Teledyne Dalsa, model M2020) outfitted with a lens (Tamron 032938) and infrared longpass filter (Midopt lp830-25.5).

### Fiber photometry

LEDs of different wavelengths provided a basis for separating ACh-dependent (465 nm) and ACh-independent (405 nm) fluorescence^7^. Blue and purple LEDs were modulated at 210 and 330 Hz, respectively, and combined through an integrated fluorescence mini-cube (FMC4, Doric). The power at the tip of the patch cable was 0.1–0.2 mW. The optical patch cable was connected to the fiber implant via a zirconia mating sleeve. Bulk fluorescent signals were acquired with a femtowatt photoreceiver (2151, Newport) and digital signal processor (Tucker-Davis Technologies RZ5D). The signal was demodulated by the lock-in amplifier implemented in the processor, sampled at 1017 Hz, and low-pass filtered with a corner frequency at 20 Hz. The optical fibers were prebleached overnight by setting both LEDs to constant illumination at a low power (<50 uW).

### Stimuli and data collection

Visual gratings were generated in MATLAB using the Psychtoolbox extension and presented via an 800 × 480 pixel display (Adafruit 2406) positioned approximately 15 cm from the left eye 45° off midline. Visual gratings were presented with a spatial frequency of 0.035 cycles per degree at three contrasts: 11%, 33%, and 100%. Gratings (2 s duration) were presented at both vertical and horizontal orientations. Spatial drift (2 Hz) was imposed along the orthogonal axis to orientation.

Auditory stimuli were either pure tones or auditory drifting gratings (i.e., ripples). Stimuli were generated with a 24-bit digital-to-analog converter (National Instruments model PXI-4461) and presented via a free-field speaker (CUI, CMS0201KLX) placed approximately 10 cm from the left (contralateral) ear canal. Free-field stimuli were calibrated using a wide-band free-field microphone (PCB Electronics, 378C01). Pure tones were low (either 6 or 6.8 kHz), mid (9.5 or 11.3 kHz), or high (13.9 or 18.5 kHz) frequencies presented at three intensities (30, 50, and 70 dB SPL). Tones were 0.4 s duration shaped with 5 ms raised cosine onset and offset ramps. Auditory gratings ranged from 2 to 45 kHz with 2 s duration (5 ms raised cosine onset and offset ramps), presented at downward and upward frequency trajectories (at –2 and +2 Hz) at three intensities (30, 50, and 70 dB SPL). The spectrum was shaped with 20 frequency carriers per octave that were sinusoidally modulated with 90% depth at one cycle per octave.

### QUANTIFICATIONS AND STATISTICAL ANALYSIS

#### Pupil and FME extraction

Pupil diameter for ACh3.0 sensor imaging experiments was extracted using DeepLabCut (version 2.1.8.2^56^). 10 frames taken from 31 mice, for a total of 310 frames, were labeled by 2-3 investigators. The four cardinal and four intercardinal compass points were marked for each pupil. Marker placement was confirmed by at least one additional investigator. Training was performed on 95% of frames. We used a ResNet-101 based neural network with default parameters for 1,030,000 training iterations. We then used a p-value cutoff of 0.9 to condition the X,Y coordinates for analysis. This network was then used to analyze videos from similar experimental settings from the ACh3.0 sensor imaging mice. We calculated pupil diameter for each frame by fitting an ellipse to the identified pupil contour points using a least-squares criterion and calculating the long axis diameter.

#### Signal Pre-processing

We first used the 465 nm ACh signal to calculate the fractional change in fluorescence DF/F_0_, where F_0_ was defined as the running median fluorescence value in a 60 s time window. DF/F_0_ traces were then low-pass filtered with a second-order zero-lag Butterworth filter, with a cut-off frequency set to 7 Hz.

#### FME bout detection and spontaneous/evoked event matching

Facial motion energy, or FME, was extracted as described previously^21^. Briefly, a rectangular region of interest (ROI) was drawn on the cheek just caudal to the vibrissae array. For each pixel in the ROI, we calculated the absolute intensity difference between consecutive frames, summing these values to generate the FME. FME was z-scored with respect to the mean and standard deviation of the entire session. For analysis, we upsampled the FME to the ACh acquisition rate (1017 Hz).

To detect discrete FME bouts corresponding to brief twitches of the face, we first smoothed the continuous FME trace with a median filter (0.5 s kernel) before calculating the global z-score. FME samples exceeding a 75^th^ percentile threshold were considered putative movement bouts. Adjacent above-threshold periods with less than 0.5 s in between were merged to avoid artificially fragmenting individual bouts of movement. Bout onsets and offsets were then identified as the first and last timepoint of each contiguous above-threshold epoch. Of the putative bouts, only those with an area under the curve (AUC) of above 0.1 (z-score; integrated from 0 to 1s post-onset) were considered in the analysis.

To analyze stimulus-evoked bouts, we defined the bout onset as the first sample after the stimulus onset that exceeded the 75^th^ percentile threshold. We used a procedure to match and balance the amplitude of stimulus evoked FME bouts to spontaneous bouts, which were gathered from sessions in which there were no stimuli. For each evoked bout, all spontaneous bouts whose amplitude AUC was within a 15% tolerance threshold of the evoked bout AUC were considered potential matches. From the candidate pool of spontaneous bout matches, an equal number were considered from -15% and +15% ranges so that spontaneous bout matches would not be systematically larger or smaller than their evoked bout counterparts. Within this balanced distribution, a spontaneous bout match was chosen for each evoked bout at random. Evoked bouts for which a balanced spontaneous amplitude match distribution could not be constructed, were excluded from the analysis.

### FIR Regression Model

We used a multiple linear regression model to quantify how well the unique stimuli, facial motion energy, and pupil diameter predicted the phasic (0-1 sec from stim onset) ACh signal. To take advantage of the differing timescales along which the predictors and the ACh signal unfold, we utilized a Finite Impulse Response (FIR) regression model, which uses time-shifted copies of the input predictors^57^ to predict the ACh response. The stimulus predictor was a delta function at stimulus onset^58^, with causal lags spanning 0 to +6 sec from onset. The FME and pupil diameter predictors were represented with both causal (0-6 sec) and acausal (−2-0 sec) lags to capture contributions of pre-trial arousal state in modulating the post-stimulus phasic ACh signal. The lagged predictor matrices were concatenated into a single design matrix *X*, and predicted the vectorized ACh signal matrix *Y* across all trials. To prevent overfitting, we used ridge regression^59^ to add a penalty term to large weights (L2 regularization, where the squared magnitude of each weight is penalized). The model’s performance was evaluated using 5-fold cross-validation. To analyze the unique contributions of each predictor (stimulus, FME, pupil diameter), we used variance partitioning. We defined each predictor’s unique R^2^ as the difference between the full-model R^2^ and the R^2^ of a partial model without that predictor.

### Statistical comparisons and repeated measures correction

All plotted data is represented at mean +/- s.e.m. or median +/- inter-quartile range for normally or non-normally distributed data, respectively. Group-level statistical comparisons utilized paired or unpaired *t-*tests or the Wilcoxon signed-rank test. Multiple comparisons correction was performed using the Benjamini-Hochberg procedure^60^.

## References

1. Yu, A.J., and Dayan, P. (2002). Acetylcholine in cortical inference. Neural Networks 15, 719–730. 10.1016/S0893-6080(02)00058-8.

2. Steriade, M. (2004). Acetylcholine systems and rhythmic activities during the waking–sleep cycle. In Progress in Brain Research (Elsevier), pp. 179–196. 10.1016/S0079-6123(03)45013-9.

3. Jiménez-Capdeville, M.E., and Dykes, R.W. (1996). Changes in cortical acetylcholine release in the rat during day and night: differences between motor and sensory areas. Neuroscience 71, 567–579. 10.1016/0306-4522(95)00439-4.

4. Disney, A.A., and Higley, M.J. (2020). Diverse Spatiotemporal Scales of Cholinergic Signaling in the Neocortex. J. Neurosci. 40, 720–725. 10.1523/JNEUROSCI.1306-19.2019.

5. McGinley, M.J., Vinck, M., Reimer, J., Batista-Brito, R., Zagha, E., Cadwell, C.R., Tolias, A.S., Cardin, J.A., and McCormick, D.A. (2015). Waking State: Rapid Variations Modulate Neural and Behavioral Responses. Neuron 87, 1143–1161. 10.1016/j.neuron.2015.09.012.

6. Sarter, M., and Lustig, C. (2020). Forebrain Cholinergic Signaling: Wired and Phasic, Not Tonic, and Causing Behavior. J. Neurosci. 40, 712–719. 10.1523/JNEUROSCI.1305-19.2019.

7. Jing, M., Li, Y., Zeng, J., Huang, P., Skirzewski, M., Kljakic, O., Peng, W., Qian, T., Tan, K., Zou, J., et al. (2020). An optimized acetylcholine sensor for monitoring in vivo cholinergic activity. Nat Methods 17, 1139–1146. 10.1038/s41592-020-0953-2.

8. Stringer, C., Pachitariu, M., Steinmetz, N., Reddy, C.B., Carandini, M., and Harris, K.D. (2019). Spontaneous behaviors drive multidimensional, brainwide activity. Science 364, eaav7893. 10.1126/science.aav7893.

9. Musall, S., Kaufman, M.T., Juavinett, A.L., Gluf, S., and Churchland, A.K. (2019). Single-trial neural dynamics are dominated by richly varied movements. Nat Neurosci 22, 1677–1686. 10.1038/s41593-019-0502-4.

10. Olsen, T., and Hasenstaub, A. (2025). Sensory origin of visually evoked activity in auditory cortex. Cell Reports 44, 116364. 10.1016/j.celrep.2025.116364.

11. Bimbard, C., Sit, T.P.H., Lebedeva, A., Reddy, C.B., Harris, K.D., and Carandini, M. (2023). Behavioral origin of sound-evoked activity in mouse visual cortex. Nat Neurosci 26, 251–258. 10.1038/s41593-022-01227-x.

12. Collins, L., Francis, J., Emanuel, B., and McCormick, D.A. (2023). Cholinergic and noradrenergic axonal activity contains a behavioral-state signal that is coordinated across the dorsal cortex. eLife 12, e81826. 10.7554/eLife.81826.

13. Chavez, C., and Zaborszky, L. (2016). Basal Forebrain Cholinergic–Auditory Cortical Network: Primary Versus Nonprimary Auditory Cortical Areas. Cereb. Cortex, bhw091. 10.1093/cercor/bhw091.

14. Moriizumi, T., and Hattori, T. (1992). Ultrastructural morphology of projections from the medial geniculate nucleus and its adjacent region to the basal ganglia. Brain Research Bulletin 29, 193–198. 10.1016/0361-9230(92)90026-T.

15. Zhu, F., Elnozahy, S., Lawlor, J., and Kuchibhotla, K.V. (2023). The cholinergic basal forebrain provides a parallel channel for state-dependent sensory signaling to auditory cortex. Nat Neurosci 26, 810–819. 10.1038/s41593-023-01289-5.

16. Guo, W., Robert, B., and Polley, D.B. (2019). The Cholinergic Basal Forebrain Links Auditory Stimuli with Delayed Reinforcement to Support Learning. Neuron 103, 1164–1177.e6. 10.1016/j.neuron.2019.06.024.

17. Robert, B., Kimchi, E.Y., Watanabe, Y., Chakoma, T., Jing, M., Li, Y., and Polley, D.B. (2021). A functional topography within the cholinergic basal forebrain for encoding sensory cues and behavioral reinforcement outcomes. eLife 10, e69514. 10.7554/eLife.69514.

18. Chernyshev, B.V., and Weinberger, N.M. (1998). Acoustic frequency tuning of neurons in the basal forebrain of the waking guinea pig. Brain Research 793, 79–94. 10.1016/S0006-8993(98)00163-2.

19. Guo, W., Robert, B., and Polley, D.B. (2019). The Cholinergic Basal Forebrain Links Auditory Stimuli with Delayed Reinforcement to Support Learning. Neuron 103, 1164–1177.e6. 10.1016/j.neuron.2019.06.024.

20. Lohani, S., Moberly, A.H., Benisty, H., Landa, B., Jing, M., Li, Y., Higley, M.J., and Cardin, J.A. (2022). Spatiotemporally heterogeneous coordination of cholinergic and neocortical activity. Nat Neurosci 25, 1706–1713. 10.1038/s41593-022-01202-6.

21. Clayton, K.K., Stecyk, K.S., Guo, A.A., Chambers, A.R., Chen, K., Hancock, K.E., and Polley, D.B. (2024). Sound elicits stereotyped facial movements that provide a sensitive index of hearing abilities in mice. Current Biology 34, 1605–1620.e5. 10.1016/j.cub.2024.02.057.

22. Williams, A.M., Angeloni, C.F., and Geffen, M.N. (2023). Sound Improves Neuronal Encoding of Visual Stimuli in Mouse Primary Visual Cortex. J. Neurosci. 43, 2885–2906. 10.1523/JNEUROSCI.2444-21.2023.

23. Zhang, G.-W., Shen, L., Zhong, W., Xiong, Y., Zhang, L.I., and Tao, H.W. (2018). Transforming Sensory Cues into Aversive Emotion via Septal-Habenular Pathway. Neuron 99, 1016–1028.e5. 10.1016/j.neuron.2018.07.023.

24. Zhang, G.-W., Sun, W.-J., Zingg, B., Shen, L., He, J., Xiong, Y., Tao, H.W., and Zhang, L.I. (2018). A Non-canonical Reticular-Limbic Central Auditory Pathway via Medial Septum Contributes to Fear Conditioning. Neuron 97, 406–417.e4. 10.1016/j.neuron.2017.12.010.

25. Mathis, A., Mamidanna, P., Cury, K.M., Abe, T., Murthy, V.N., Mathis, M.W., and Bethge, M. (2018). DeepLabCut: markerless pose estimation of user-defined body parts with deep learning. Nat Neurosci 21, 1281–1289. 10.1038/s41593-018-0209-y.

26. Vinck, M., Batista-Brito, R., Knoblich, U., and Cardin, J.A. (2015). Arousal and Locomotion Make Distinct Contributions to Cortical Activity Patterns and Visual Encoding. Neuron 86, 740–754. 10.1016/j.neuron.2015.03.028.

27. Asokan, M.M., Watanabe, Y., Kimchi, E.Y., and Polley, D.B. (2023). Potentiation of cholinergic and corticofugal inputs to the lateral amygdala in threat learning. Cell Reports 42, 113167. 10.1016/j.celrep.2023.113167.

28. Woolley, S.M.N., Fremouw, T.E., Hsu, A., and Theunissen, F.E. (2005). Tuning for spectro-temporal modulations as a mechanism for auditory discrimination of natural sounds. Nat Neurosci 8, 1371–1379. 10.1038/nn1536.

29. Zimmer-Harwood, P., Picard, S., King, A.J., and Dahmen, J.C. (2026). Auditory cortex distinguishes between spontaneous and sound-evoked movements. J. Neurosci., e1874252026. 10.1523/JNEUROSCI.1874-25.2026.

30. Zou, J., Willem De Gee, J., Mridha, Z., Trinh, S., Erskine, A., Jing, M., Yao, J., Walker, S., Li, Y., McGinley, M., et al. (2024). Goal-directed motor actions drive acetylcholine dynamics in sensory cortex. Preprint, 10.7554/eLife.96931.1 http://doi.org/10.7554/eLife.96931.1.

31. Mouland, J.W., Tamayo, E., Ebrahimi, A.S., Williams, C., Fleming, W., Watson, A., Hogan, M.P., Lucas, R.J., Storchi, R., and Brown, T.M. (2025). A lateral hypothalamic region supporting diverse visual processing and modulation of visually-guided behaviour. Nat Commun 16, 9917. 10.1038/s41467-025-64864-3.

32. Martianova, E., Sadretdinova, R., Pageau, A., Pausic, N., Gentiletti, T.D., Leblanc, D., Rivera, A.M., Labonté, B., and Proulx, C.D. (2023). Hypothalamic neuronal outputs transmit sensorimotor signals at the onset of locomotor initiation. iScience 26, 108328. 10.1016/j.isci.2023.108328.

33. Hayat, H., Regev, N., Matosevich, N., Sales, A., Paredes-Rodriguez, E., Krom, A.J., Bergman, L., Li, Y., Lavigne, M., Kremer, E.J., et al. (2020). Locus coeruleus norepinephrine activity mediates sensory-evoked awakenings from sleep. Sci. Adv. 6, eaaz4232. 10.1126/sciadv.aaz4232.

34. Ranade, S.P., and Mainen, Z.F. (2009). Transient Firing of Dorsal Raphe Neurons Encodes Diverse and Specific Sensory, Motor, and Reward Events. Journal of Neurophysiology 102, 3026–3037. 10.1152/jn.00507.2009.

35. Mutlu, A.K., Serneels, B., Wiest, C., Trinh, A.-T., Bardenhewer, R., Palumbo, F., Frisvold, O.B., Aukrust, I.K.F., Ostenrath, A.M., and Yaksi, E. (2025). Topographically organized dorsal raphe activity modulates forebrain sensory-motor computations and adaptive behaviors. Preprint at Neuroscience, http://doi.org/10.1101/2025.03.06.641626 10.1101/2025.03.06.641626.

36. Hulsey, D., Zumwalt, K., Mazzucato, L., McCormick, D.A., and Jaramillo, S. (2024). Decision-making dynamics are predicted by arousal and uninstructed movements. Cell Reports 43, 113709. 10.1016/j.celrep.2024.113709.

37. Edeline, J., Dutrieux, G., Manunta, Y., and Hennevin, E. (2001). Diversity of receptive field changes in auditory cortex during natural sleep. Eur J of Neuroscience 14, 1865–1880. 10.1046/j.0953-816x.2001.01821.x.

38. Wang, Y., You, L., Tan, K., Li, M., Zou, J., Zhao, Z., Hu, W., Li, T., Xie, F., Li, C., et al. (2023). A common thalamic hub for general and defensive arousal control. Neuron, S0896627323005408. 10.1016/j.neuron.2023.07.007.

39. Boccalaro, I.L., Aime, M., Aellen, F.M., Rusterholz, T., Borsa, M., Bozic, I., Sattin, A., Fellin, T., Herrera, C.G., Tzovara, A., et al. (2025). A role for the thalamus in danger evoked awakening during sleep. Nat Commun 16, 7049. 10.1038/s41467-025-62265-0.

40. Gielow, M.R., and Zaborszky, L. (2017). The Input-Output Relationship of the Cholinergic Basal Forebrain. Cell Reports 18, 1817–1830. 10.1016/j.celrep.2017.01.060.

41. Kim, J.-H., Jung, A.-H., Jeong, D., Choi, I., Kim, K., Shin, S., Kim, S.J., and Lee, S.-H. (2016). Selectivity of Neuromodulatory Projections from the Basal Forebrain and Locus Ceruleus to Primary Sensory Cortices. J. Neurosci. 36, 5314–5327. 10.1523/JNEUROSCI.4333-15.2016.

42. Reimer, J., McGinley, M.J., Liu, Y., Rodenkirch, C., Wang, Q., McCormick, D.A., and Tolias, A.S. (2016). Pupil fluctuations track rapid changes in adrenergic and cholinergic activity in cortex. Nat Commun 7, 13289. 10.1038/ncomms13289.

43. Grujic, N., Polania, R., and Burdakov, D. (2024). Neurobehavioral meaning of pupil size. Neuron 112, 3381–3395. 10.1016/j.neuron.2024.05.029.

44. Bertero, A., and Apicella, A.J. (2026). A non-canonical cholinergic pathway from the dorsal tail of the striatum to the auditory cortex. Nat Commun. 10.1038/s41467-026-72939-y.

45. Schlingloff, D., Szabó, í., Gulyás, É., Király, B., Kispál, R., Stephenson-Jones, M., and Hangya, B. (2026). Most Ventral Pallidal Cholinergic Neurons Are Bursting Basal Forebrain Cholinergic Neurons with Mesocorticolimbic Connectivity. J. Neurosci. 46, e0415252026. 10.1523/JNEUROSCI.0415-25.2026.

46. Granger, A.J., Wang, W., Robertson, K., El-Rifai, M., Zanello, A.F., Bistrong, K., Saunders, A., Chow, B.W., Nuñez, V., Turrero García, M., et al. (2020). Cortical ChAT+ neurons co-transmit acetylcholine and GABA in a target-and brain-region-specific manner. eLife 9, e57749. 10.7554/eLife.57749.

47. Meyer, A.F., Poort, J., O’Keefe, J., Sahani, M., and Linden, J.F. (2018). A Head-Mounted Camera System Integrates Detailed Behavioral Monitoring with Multichannel Electrophysiology in Freely Moving Mice. Neuron 100, 46–60.e7. 10.1016/j.neuron.2018.09.020.

48. Xu, M., Chung, S., Zhang, S., Zhong, P., Ma, C., Chang, W.-C., Weissbourd, B., Sakai, N., Luo, L., Nishino, S., et al. (2015). Basal forebrain circuit for sleep-wake control. Nat Neurosci 18, 1641–1647. 10.1038/nn.4143.

49. Picciotto, M.R., Higley, M.J., and Mineur, Y.S. (2012). Acetylcholine as a Neuromodulator: Cholinergic Signaling Shapes Nervous System Function and Behavior. Neuron 76, 116–129. 10.1016/j.neuron.2012.08.036.

50. Vattino, L.G., Clayton, K.K., Hackett, T.A., Polley, D.B., and Takesian, A.E. (2026). Layer 6 is a hub for cholinergic modulation in the mouse auditory cortex. Cerebral Cortex 36, bhaf338. 10.1093/cercor/bhaf338.

51. Pinto, L., Goard, M.J., Estandian, D., Xu, M., Kwan, A.C., Lee, S.-H., Harrison, T.C., Feng, G., and Dan, Y. (2013). Fast modulation of visual perception by basal forebrain cholinergic neurons. Nat Neurosci 16, 1857–1863. 10.1038/nn.3552.

52. Nodal, F.R., Leach, N.D., Keating, P., Dahmen, J.C., Zhao, D., King, A.J., and Bajo, V.M. (2024). Neural processing in the primary auditory cortex following cholinergic lesions of the basal forebrain in ferrets. Hearing Research 447, 109025. 10.1016/j.heares.2024.109025.

53. Záborszky, L., Gombkoto, P., Varsanyi, P., Gielow, M.R., Poe, G., Role, L.W., Ananth, M., Rajebhosale, P., Talmage, D.A., Hasselmo, M.E., et al. (2018). Specific Basal Forebrain– Cortical Cholinergic Circuits Coordinate Cognitive Operations. J. Neurosci. 38, 9446–9458. 10.1523/JNEUROSCI.1676-18.2018.

54. Parikh, V., Kozak, R., Martinez, V., and Sarter, M. (2007). Prefrontal Acetylcholine Release Controls Cue Detection on Multiple Timescales. Neuron 56, 141–154. 10.1016/j.neuron.2007.08.025.

55. Minces, V., Pinto, L., Dan, Y., and Chiba, A.A. (2017). Cholinergic shaping of neural correlations. Proc. Natl. Acad. Sci. U.S.A. 114, 5725–5730. 10.1073/pnas.1621493114.

56. Nath, T., Mathis, A., Chen, A.C., Patel, A., Bethge, M., and Mathis, M.W. (2019). Using DeepLabCut for 3D markerless pose estimation across species and behaviors. Nat Protoc 14, 2152–2176. 10.1038/s41596-019-0176-0.

57. Dale, A.M. (1999). Optimal experimental design for event-related fMRI. Hum. Brain Mapp. 8, 109–114. 10.1002/(SICI)1097-0193(1999)8:2/3%3C109::AID-HBM7%3E3.0.CO;2-W.

58. Simoncelli, E.P., Paninski, L., Pillow, J., and Schwartz, O. Characterization of Neural Responses with Stochastic Stimuli.

59. Hoerl, A.E., and Kennard, R.W. Ridge Regression: Biased Estimation for Nonorthogonal Problems.

60. Benjamini, Y., and Hochberg, Y. (1995). Controlling the False Discovery Rate: A Practical and Powerful Approach to Multiple Testing. Journal of the Royal Statistical Society Series B: Statistical Methodology 57, 289–300. 10.1111/j.2517-6161.1995.tb02031.x.

