## Supplemental Information for "Spontaneous and stimulus-driven arousal produce distinct acetylcholine dynamics across sensory and prefrontal cortex"

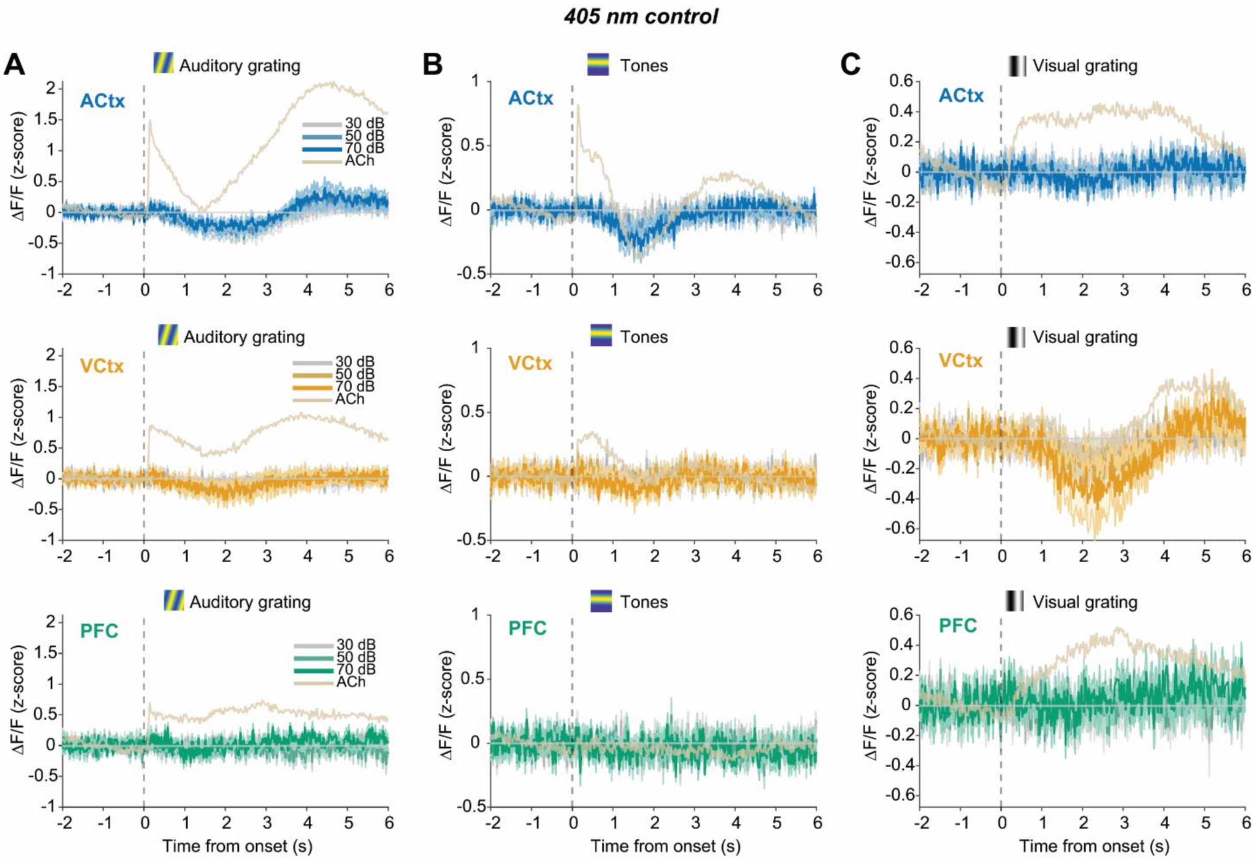


**FIGURE S1.** (**A**) Auditory grating-evoked DF/F traces (z-scored) at different intensities, across ACtx, VCtx, and PFC (top to bottom), in the 405 nm isosbestic control condition. Beige trace in each plot shows the corresponding ACh (465 nm) response in each cortical region, pooled across intensities, for comparison. (**B-C**) Same as **A**, but for tones (**B**) and visual gratings (**C**).

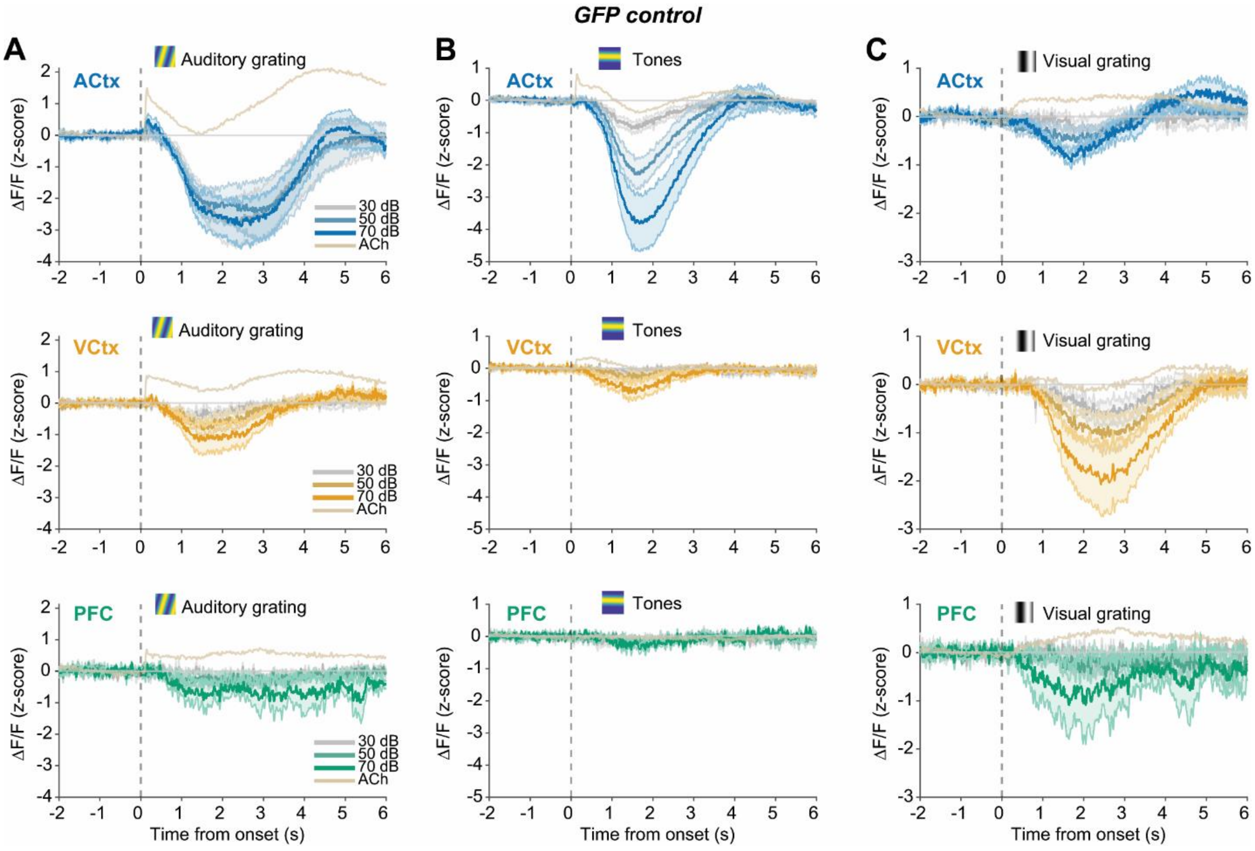


**FIGURE S2.** (**A**) Auditory grating-evoked DF/F traces (z-scored) at different intensities, across ACtx, VCtx, and PFC (top to bottom), in mice expressing the fluorophore GFP in cortex, rather than the ACh indicator. Beige trace in each plot shows the corresponding ACh (465 nm) response in each cortical region, pooled across intensities, for comparison. (**B-C**) Same as **A**, but for tones (**B**) and visual gratings (**C**).

| Signal | Stimulus | Region | Intensity | n_mice | Mean_AUC | SEM_AUC | p_raw_Wilcoxon |
|---|---|---|---|---|---|---|---|
| ACh | Aud. Ripple | AC | 30 dB | 11 | 0.553 | 0.1994 | 0.002 |
| ACh | Aud. Ripple | AC | 50 dB | 11 | 0.5739 | 0.2357 | 0.002 |
| ACh | Aud. Ripple | AC | 70 dB | 11 | 0.9538 | 0.2711 | 0.002 |
| ACh | Aud. Ripple | VC | 30 dB | 7 | 0.4931 | 0.1293 | 0.0156 |
| ACh | Aud. Ripple | VC | 50 dB | 7 | 0.6589 | 0.1374 | 0.0156 |
| ACh | Aud. Ripple | VC | 70 dB | 7 | 0.7577 | 0.1603 | 0.0156 |
| ACh | Aud. Ripple | PFC | 30 dB | 5 | 0.0747 | 0.1104 | 1 |
| ACh | Aud. Ripple | PFC | 50 dB | 5 | 0.3184 | 0.1725 | 0.1875 |
| ACh | Aud. Ripple | PFC | 70 dB | 5 | 0.8759 | 0.2061 | 0.0625 |
| ACh | Tones | AC | 30 dB | 11 | 0.2507 | 0.0862 | 0.0098 |
| ACh | Tones | AC | 50 dB | 11 | 0.3174 | 0.1159 | 0.0098 |
| ACh | Tones | AC | 70 dB | 11 | 0.4763 | 0.1642 | 0.0068 |
| ACh | Tones | VC | 30 dB | 7 | 0.0214 | 0.0307 | 0.8125 |
| ACh | Tones | VC | 50 dB | 7 | 0.2821 | 0.0872 | 0.0156 |
| ACh | Tones | VC | 70 dB | 7 | 0.4038 | 0.111 | 0.0156 |
| ACh | Tones | PFC | 30 dB | 5 | -0.0581 | 0.05 | 0.3125 |
| ACh | Tones | PFC | 50 dB | 5 | -0.063 | 0.0721 | 0.625 |
| ACh | Tones | PFC | 70 dB | 5 | 0.039 | 0.0359 | 0.4375 |
| ACh | Vis. Grating | AC | 0.11 | 11 | 0.0638 | 0.0571 | 0.5771 |
| ACh | Vis. Grating | AC | 0.33 | 11 | 0.01 | 0.0548 | 0.5771 |
| ACh | Vis. Grating | AC | 1 | 11 | 0.6029 | 0.1637 | 0.001 |
| ACh | Vis. Grating | VC | 0.11 | 7 | -0.101 | 0.0699 | 0.1562 |
| ACh | Vis. Grating | VC | 0.33 | 7 | 0.0365 | 0.0659 | 0.6875 |
| ACh | Vis. Grating | VC | 1 | 7 | 0.2271 | 0.0588 | 0.0312 |
| ACh | Vis. Grating | PFC | 0.11 | 5 | -0.1016 | 0.0683 | 0.3125 |
| ACh | Vis. Grating | PFC | 0.33 | 5 | -0.028 | 0.0216 | 0.4375 |
| ACh | Vis. Grating | PFC | 1 | 5 | 0.3771 | 0.1168 | 0.0625 |

| | | | | | | | |
|---|---|---|---|---|---|---|---|
| GFP | Aud. Ripple | AC | 30 dB | 6 | -0.2095 | 0.1164 | 0.1562 |
| GFP | Aud. Ripple | AC | 50 dB | 6 | -0.2314 | 0.1007 | 0.0938 |
| GFP | Aud. Ripple | AC | 70 dB | 6 | -0.1262 | 0.1251 | 0.5625 |
| GFP | Aud. Ripple | VC | 30 dB | 6 | -0.1292 | 0.0395 | 0.0312 |
| GFP | Aud. Ripple | VC | 50 dB | 6 | -0.1416 | 0.062 | 0.1562 |
| GFP | Aud. Ripple | VC | 70 dB | 6 | -0.1672 | 0.0564 | 0.0312 |
| GFP | Aud. Ripple | PFC | 30 dB | 3 | -0.0369 | 0.0227 | 0.25 |
| GFP | Aud. Ripple | PFC | 50 dB | 3 | -0.1007 | 0.0642 | 0.25 |
| GFP | Aud. Ripple | PFC | 70 dB | 3 | -0.1695 | 0.0798 | 0.25 |
| GFP | Tones | AC | 30 dB | 6 | -0.0976 | 0.0456 | 0.1562 |
| GFP | Tones | AC | 50 dB | 6 | -0.2162 | 0.1044 | 0.1562 |
| GFP | Tones | AC | 70 dB | 6 | -0.4568 | 0.1132 | 0.0312 |
| GFP | Tones | VC | 30 dB | 6 | -0.0379 | 0.0227 | 0.1562 |
| GFP | Tones | VC | 50 dB | 6 | -0.0644 | 0.0427 | 0.2188 |
| GFP | Tones | VC | 70 dB | 6 | -0.2127 | 0.104 | 0.0625 |
| GFP | Tones | PFC | 30 dB | 3 | -0.0982 | 0.0873 | 0.5 |
| GFP | Tones | PFC | 50 dB | 3 | -0.0095 | 0.0282 | 0.75 |
| GFP | Tones | PFC | 70 dB | 3 | -0.0194 | 0.0174 | 0.5 |
| GFP | Vis. Grating | AC | 0.11 | 6 | -0.0315 | 0.0841 | 1 |
| GFP | Vis. Grating | AC | 0.33 | 6 | -0.244 | 0.0987 | 0.0312 |
| GFP | Vis. Grating | AC | 1 | 6 | -0.1551 | 0.0692 | 0.1562 |
| GFP | Vis. Grating | VC | 0.11 | 6 | -0.0946 | 0.0552 | 0.3125 |
| GFP | Vis. Grating | VC | 0.33 | 6 | -0.0321 | 0.0706 | 0.4375 |
| GFP | Vis. Grating | VC | 1 | 6 | -0.0647 | 0.0305 | 0.1562 |
| GFP | Vis. Grating | PFC | 0.11 | 3 | 0.0253 | 0.0356 | 0.5 |
| GFP | Vis. Grating | PFC | 0.33 | 3 | -0.0803 | 0.0404 | 0.25 |
| GFP | Vis. Grating | PFC | 1 | 3 | -0.2199 | 0.2227 | 0.75 |

| | | | | | | | |
|---|---|---|---|---|---|---|---|
| 405nm | Aud. Ripple | AC | 30 dB | 10 | -0.0084 | 0.0198 | 0.9219 |
| 405nm | Aud. Ripple | AC | 50 dB | 10 | -0.0517 | 0.0304 | 0.1309 |
| 405nm | Aud. Ripple | AC | 70 dB | 10 | 0.0054 | 0.0399 | 0.4922 |
| 405nm | Aud. Ripple | VC | 30 dB | 7 | 0.0003 | 0.0072 | 0.9375 |
| 405nm | Aud. Ripple | VC | 50 dB | 7 | -0.0473 | 0.0159 | 0.0312 |
| 405nm | Aud. Ripple | VC | 70 dB | 7 | -0.0231 | 0.0284 | 0.8125 |
| 405nm | Aud. Ripple | PFC | 30 dB | 4 | -0.0059 | 0.0055 | 0.625 |
| 405nm | Aud. Ripple | PFC | 50 dB | 4 | -0.022 | 0.0112 | 0.125 |
| 405nm | Aud. Ripple | PFC | 70 dB | 4 | 0.0301 | 0.0162 | 0.125 |
| 405nm | Tones | AC | 30 dB | 10 | 0.0072 | 0.0115 | 0.4316 |
| 405nm | Tones | AC | 50 dB | 10 | -0.0255 | 0.0246 | 0.625 |
| 405nm | Tones | AC | 70 dB | 10 | -0.0207 | 0.0229 | 0.625 |
| 405nm | Tones | VC | 30 dB | 7 | 0.0049 | 0.0066 | 0.4688 |
| 405nm | Tones | VC | 50 dB | 7 | 0.02 | 0.0128 | 0.1562 |
| 405nm | Tones | VC | 70 dB | 7 | -0.0301 | 0.0213 | 0.1562 |
| 405nm | Tones | PFC | 30 dB | 4 | 0.0021 | 0.0251 | 0.875 |
| 405nm | Tones | PFC | 50 dB | 4 | -0.0104 | 0.0266 | 0.875 |
| 405nm | Tones | PFC | 70 dB | 4 | -0.0256 | 0.0081 | 0.125 |
| 405nm | Vis. Grating | AC | 0.11 | 10 | 0.022 | 0.0276 | 0.7695 |
| 405nm | Vis. Grating | AC | 0.33 | 10 | -0.0009 | 0.0182 | 1 |
| 405nm | Vis. Grating | AC | 1 | 10 | -0.0024 | 0.0137 | 0.6953 |
| 405nm | Vis. Grating | VC | 0.11 | 7 | -0.0198 | 0.0183 | 0.4688 |
| 405nm | Vis. Grating | VC | 0.33 | 7 | -0.0107 | 0.0219 | 0.375 |
| 405nm | Vis. Grating | VC | 1 | 7 | -0.0105 | 0.0159 | 0.6875 |
| 405nm | Vis. Grating | PFC | 0.11 | 4 | 0.0057 | 0.0114 | 0.875 |
| 405nm | Vis. Grating | PFC | 0.33 | 4 | 0.0311 | 0.0272 | 0.375 |
| 405nm | Vis. Grating | PFC | 1 | 4 | 0.0256 | 0.0173 | 0.25 |

**TABLE S1**. Results from one-sided *t*-test comparing the distribution of mean AUC values (from 0-1 s post stimulus onset) of ACh, isosbestic control (405nm), or GFP z-scored DF/F in each cortical region, for each stimulus type.
